# Novel Class B2 and C β-lactamases harboured by *Pseudomonas* spp. wastewater isolates

**DOI:** 10.64898/2026.02.12.705516

**Authors:** Alexander D. H. Kingdon, Ellie Allman, Anya Breen, Kara D’Arcy, Claudia McKeown, Fabrice E. Graf, Adam P. Roberts

## Abstract

**Introduction:** Antimicrobial resistance has existed in the environment long before its rapid emergence and detection in clinically relevant pathogens. Studying the resistance of environmental bacterial strains may allow novel resistance mechanisms to be identified before they appear in pathogenic strains.

**Gap Statement:** Searching for antimicrobial resistance genes in environmental bacteria represents an understudied research area compared to resistance within clinically relevant pathogens.

**Aim:** To evaluate resistance genes present within environmental non-aeruginosa *Pseudomonas* spp. isolates.

**Methodology:** We screened a set of bacterial isolates from untreated wastewater from Liverpool, UK, for the presence of extended spectrum β-lactamases and carbapenemases. A sub-set of three resistant *Pseudomonas* spp. isolates were selected for whole-genome sequencing. We performed minimum inhibitory concentration assays against several β-lactams, and ectopic expression of four novel resistance genes within *Escherichia coli*.

**Results:** Here, we report the discovery of novel class C β-lactamase genes *bla_PFL7_, bla_PFL8_* and *bla_PFL9_*, as well as a novel subclass B2 metallo-β-lactamase *bla_PFM5_* present within these strains. The class C genes encoded proteins with between 61-71% amino acid identity to the closest known match, *bla_PFL-1_*. These novel β-lactamases degraded the cephalosporin nitrocefin and confer piperacillin and ceftazidime resistance to susceptible *Escherichia coli* when ectopically expressed. The β-lactamase inhibitor tazobactam was effective at inhibiting these enzymes. The sub-class B2 metallo-β-lactamase had 88% amino acid identity to its closet match *bla_PFM-1_* and conferred carbapenem resistance to susceptible *E. coli*. The β-lactamase inhibitors relebactam, vaborbactam, xeruborbactam and captopril had no impact on the carbapenem resistance phenotype. Analogues of all these novel genes (>95% nucleotide sequence identity) were identified within publicly available whole-genome sequencing data, suggesting they are found sporadically.

**Conclusion:** Our analysis adds to the growing number of β-lactamase genes found from environmental *Pseudomonas* spp. and suggests that continued surveillance of this environmental reservoir for novel, clinically relevant, β-lactamase genes is warranted.

## Introduction

Drug resistant *Pseudomonas aeruginosa* was associated with between 234,000-457,000 deaths in 2019, being directly attributable to 53,000-127,000 of these deaths [1]. Approximately seventy-four percent of these drug-resistant *P. aeruginosa* were resistant to β-lactams, either penicillins, cephalosporins, carbapenems, or a combination of all three, attributable to 38,000-98,100 deaths [1]. Rates of resistance are continually increasing, and resistance has been found for all new β-lactams, including more recently approved pairs of β-lactam - β-lactamase inhibitor, such as cefepime-enmetazobactam and ceftazidime-avibactam [2–4]. Growing resistance is especially problematic as β-lactams alone and in combination with β-lactamase inhibitors, are typically used as both first-and second-line treatment options for *Pseudomonas aeruginosa* infections [5]. Gaining a greater understanding of the resistance mechanisms which are already present in the environment can help shape future β-lactam development, avoiding known resistance mechanisms.

Wastewater has been shown to contain many antimicrobial resistance genes (ARGs), and wastewater treatment plants can increase presence of ARGs in downstream rivers [6,7]. Monitoring of wastewater has been undertaken for the presence of several infectious agents and has driven the development of wastewater-based epidemiology [8,9]. However, monitoring for the presence of ARGs within wastewater is typically limited to academic studies or prioritised by clinical relevance of the bacterial host. Resistance mechanisms found in environmental species have historically been underexplored [10]. Hence, investigation of environmental ARGs is necessary to uncover possible future threats to antibiotic efficacy.

Multiple gene databases have been compiled to allow analysis of a genome of interest against comprehensive lists of ARGs [11]. Several of these databases exist as webservers which aim to be readily searchable, such as ResFinder, ARG-ANNOT and CARD [12–14]. The latter also provides a repository of published experimentally characterised ARGs [12]. While these databases cover a broad range of ARGs, they can lack comprehensive coverage of all sub-types. More specialist databases exist, such as the β-lactamase database (BLDB), which is a more complete list of β-lactamase genes but its webserver is only searchable using single DNA or protein sequences [15]. These databases have a bias towards resistance genes found in clinical settings, hence, an over reliance on them when screening environmental strains for ARGs may lead to an underestimation of antimicrobial resistance (AMR).

Within *P. aeruginosa*, β-lactam resistance can be caused by a combination of several factors. These include limiting permeability of drugs through the membrane, upregulation of efflux pumps, or production of β-lactamases. The reduction in membrane permeability is caused by loss or mutations of, for example, the OprD outer membrane porin [16]. Several efflux pumps, such as MexAB-OprM can confer drug, including antibiotic, resistance, especially if mutations lead to higher expression [17–19]. β-lactamases directly inactivate the drug by enzymatic cleavage of the beta-lactam. *Pseudomonas* spp. has been shown to harbour all four classes of β-lactamases, A-D. Classes A, C and D are serine β-lactamases, requiring an active site serine residue for functionality, while class B are metallo-enzymes requiring one or two zinc residues for catalytic activity. Class B can degrade the widest range of β-lactams and are not inhibited by any clinically approved β-lactamase inhibitors [20]. Whereas the serine β-lactamases have more specific activity and are targeted by β-lactamases inhibitors.

The most common β-lactamase in *P. aeruginosa*, is the chromosomally encoded class C *bla_PDC_* (*Pseudomonas*-derived cephalosporinases), having greater than 600 protein-level variants [21,22]. In addition, class B metallo-β-lactamases have been acquired by clinical *P. aeruginosa* isolates, including *bla_VIM-1_* and *bla_VIM-2_* [23,24], and several *bla_OXA_* class D β-lactamase variants [25,26]. In contrast, there have been limited studies on non-aeruginosa *Pseudomonas* β-lactamases, however, all four classes of β-lactamases have been shown to be present: Class A *bla_LUT_* in *P. luteola* [27], Class B1 *bla_PST1-2_* in *P. stutzeri* [28], Class B2 *bla_PFM_ Pseudomonas sp.* [29,30], Class B3 *bla_PAM_* and *bla_POM_* in *P. alcaligenes* and *P. otitidis* [31,32], Class C *bla_PFL_* in *P. fluorescens* [33], and Class D *bla_OXA_* in *P. putida* [34]. For Class C β-lactamases, only eight genes have been detected in non-aeruginosa *Pseudomonas* spp. strains. Two have been published with phenotypic resistance data, *bla_PFL-P1_* and *bla_PRC-1_* [33,35], both showing resistance to penicillins and cephalosporins. For Class B2 β-lactamases, four *bla_PFM_* genes have been found in non-aeruginosa *Pseudomonas* spp. [29,30]. Only *bla_PRC-1_* and *bla_PFM1-4_* are present in CARD, while none of these twelve genes are in ARG-ANNOT or ResFinder. In contrast, all three databases contain multiple *bla_PDC_* genes, 14 in ARG-ANNOT, 475 in CARD, and 4 in ResFinder. Hence, there is a large disparity between our knowledge of *P. aeruginosa* and other *Pseudomonas* β-lactam resistance genes.

As part of the Swab and Send initiative within the Infection Innovation Consortium (iiCON) at the Liverpool School of Tropical Medicine [36], we have accumulated a large microbial isolate library (∼80k isolates), which can be leveraged to identify ARGs from various environments. In this study, we screened a subset of isolates from sewage and associated wastewater [37]. We have identified four putative β-lactamase genes, three class C and one subclass B2, which were not present in any AMR gene databases. We have shown these putative β-lactamases have catalytic activity, despite relatively low amino acid sequence identity to known *Pseudomonas* β-lactamases.

## Methods

### Isolation of microbes from sewage

Sewage was collected from the external outflow pipes of Liverpool School of Tropical Medicine using an automated sampling device, storing up to one litre at a time, as previously described [37]. This sewage was swabbed after collection and swabs stored in Amies Charcoal agar (MWE) at 4 °C, prior to processing. Swabs were spread onto brain-heart infusion (BHI) agar (ThermoFisher) and incubated at room temperature for two days. Colonies with distinct morphologies were selected from each plate and transferred to individual wells containing 100 µL BHI broth (ThermoFisher) in 96-well plates. These plates were incubated statically at room temperature for two days, 100 µL BHI containing 40% glycerol (v/v) (Fisher) was then added to each well and plates were stored at –70 °C.

### Growth Conditions

Bacterial strains are listed in **Table 1 & 2**. All bacterial strains were grown using BHI broth or agar. All cultures and plates were incubated at room temperature. Broth cultures were prepared by inoculating with a single colony and grown shaking at 160 rpm at 25 °C. Long-term strain storage was undertaken using BHI containing 20% glycerol, stored at –70 °C.

**Table 1:** Species identification and β-lactam resistance of tested *Pseudomonas* spp. Strains.

| Strain ID | Isolation Date | Species Finder | gyrB sequence group [41] | MALDI-TOF | ANI Comparison (coverage) | Piperacillin MIC ($\mu\text{g/mL}$) | Ceftazidime MIC ($\mu\text{g/mL}$) | Meropenem MIC ($\mu\text{g/mL}$) | Imipenem MIC ($\mu\text{g/mL}$) |
| --- | --- | --- | --- | --- | --- | --- | --- | --- | --- |
| C001-2B | 17/03/21 | Failed (<98%) | *P. protegens* | *P. koreensis* (1.87 = low confidence) | 89.30 (85.82) to *P. piscis* | 8 | 2-4 | 16 | 8 |
| C001-7H | 17/03/21 | Failed (<98%) | *P. protegens* | *P. protegens* (1.99 = low confidence) | 98.11 (83.86) to *P. saponiphila* | 32 | 16 | 64 | <2 |
| C006-8D | 18/11/22 | Failed (<98%) | *P. fluorescens* | *P. extremorientalis* (2.00 = high confidence) | 95.97 (83.26) to *P. palleroniana* | 4 | 4 | 32 | 16-32 |

**Table 2:** *E. coli* transformants with pEB1 containing cloned novel β-lactamase genes. N=3. Tazobactam and Xeruborbactam concentration fixed at 4 µg/mL. Vaborbactam fixed at 8 µg/mL. Captopril fixed at 128 µg/mL.

| Strain + Gene | Piperacillin MIC ($\mu\text{g/mL}$) | Piperacillin + Tazobactam MIC ($\mu\text{g/mL}$) | Ceftazidime MIC ($\mu\text{g/mL}$) | Meropenem MIC ($\mu\text{g/mL}$) | Meropenem + Vaborbactam MIC ($\mu\text{g/mL}$) | Meropenem + Xeruborbactam MIC ($\mu\text{g/mL}$) | Meropenem + Captopril MIC ($\mu\text{g/mL}$) | Imipenem MIC ($\mu\text{g/mL}$) | Imipenem + Relebactam MIC ($\mu\text{g/mL}$) |
| --- | --- | --- | --- | --- | --- | --- | --- | --- | --- |
| *E. coli* + *bla*<sub>PFL7</sub> (from C001-2B) | 32-64 | 1 | 4 | <0.125 | N/A | N/A | N/A | 0.125 | N/A |
| *E. coli* + *bla*<sub>PFL8</sub> (from C001-7H) | 64 | 2 | 16 | <0.125 | N/A | N/A | N/A | 0.125 | N/A |
| *E. coli* + *bla*<sub>PFL9</sub> (from C006-8D) | 32 | 1 | 4 | <0.125 | N/A | N/A | N/A | 0.125 | N/A |
| *E. coli* + *bla*<sub>PFM5</sub> (from C006-8D) | 0.5 | N/A | <0.125 | 4 | 4 | 4 | 4 | 8 | 8 |
| *E. coli* | 0.5 | 0.5 | <0.125 | <0.125 | <0.125 | <0.125 | <0.125 | 0.125 | 0.125 |
| *E. coli* + GFP | 0.5 | 0.5 | <0.125 | <0.125 | <0.125 | <0.125 | <0.125 | 0.125 | 0.125 |
| EUCAST Breakpoints (Enterobacterales) | >8 | >8 | >4 | >2-8 | >8 | N/A | N/A | >4 | >2 |

### Antibiotic Resistance Screening

β-lactam resistant isolates were identified by stamping out frozen glycerol stocks directly onto BHI + meropenem (10 µg/mL, Sigma Aldrich) agar and CHROMagar^TM^ ESBL agar (CHROMagar). The plates were incubated at room temperature for two days. Any colonies which grew were sub-cultured onto identical plates (containing meropenem or ESBL supplement) to confirm resistance. Identified *Pseudomonas* isolates which could grow on both meropenem and ESBL supplement were selected for identification and whole genome sequencing (WGS).

### Whole Genome Sequencing

DNA extraction, WGS and genome assembly and annotation were performed by MicrobesNG. Strains were streaked out for single colonies; an overnight culture of 10 mL BHI broth was inoculated with one colony and allowed to grow until mid-log (OD_600_= 0.5-1.2) was reached. The cells were pelleted, washed with PBS and transferred to DNA/RNA Shield (Zymo Research), prior to shipping to MicrobesNG. DNA extraction was undertaken by first lysing the cells through addition of 120 μL of TE buffer containing lysozyme (MPBio), metapolyzyme (Sigma Aldrich) and RNase A (ITW Reagents). This was incubated at 37 °C for 25 minutes, before addition of proteinase K (VWR chemicals, 0.1 mg/mL) and SDS (0.5% v/v) and incubation at 65°C for 5 minutes. The genomic DNA was purified using SPRI beads (equal volume) and resuspended in EB buffer (10 mM Tris-HCl, pH 8.0). The library preparation was undertaken using a Nextera XT library prep kit (Illumina), following the manufacturer’s protocol except for doubling of the input DNA and an increase in the elongation time of the PCR to 45 seconds. A short read 250 bp paired end protocol for sequencing was undertaken on an Illumina NovaSeq6000.

Raw reads were trimmed using Trimmomatic v0.3 [38], with a sliding window cut-off of Q15. A minimum mean genome coverage of 30x was used as a quality cut-off. De novo genome assembly was performed using SPAdes v3.7 [39], and contigs were annotated using Prokka v1.11 [40].

### Species Identification

The *gyrB* gene sequences were taken from the assembled genomes of our sequenced strains and compared to other *gyrB* genes from publicly available genomes of several *Pseudomonas* spp. (**Supplementary Table 1**) [41,42]. These were partially grouped according to the multilocus sequence analysis inferred phylogeny of the *P. fluorescens* complex [41].

Species identification was also undertaken using a Bruker MALDI Biotyper^®^ sirius system. A sample from a single colony of each isolate was loaded onto an MBT Biotarget plate and prepared following the extended direct transfer technique advised by Bruker. Briefly, this involved addition of 70% formic acid (1 µL, VWR chemicals) and subsequent drying, prior to HCCA matrix addition (1 µL, Bruker). All runs were validated using Bruker’s bacterial test standard. All samples’ spectra were matched against the MBT Compass library to provide species ID.

### β-lactamase gene identification

Assembled genomes for the investigated *Pseudomonas* spp. strains were uploaded to the ResFinder v4.4 [14,43], CARD [12], and ARG-ANNOT V6 [13] webservers to search for ARG presence. For CARD’s Resistance Gene Identifier, a DNA sequence input was used to search for perfect, strict and loose hits. For ResFinder, the default search parameters of 90% nucleotide identity across 60% minimum length were initially used, but these were reduced to 30% ID, 40% length to detect a match. For matches which were identified in ARG-ANNOT, the genome annotation labelling was consistently ‘ampC’, for the genes present in our strains. The genomes which gave no matches in all three databases were manually searched for ‘ampC’ in their GenBank files. Once a nucleotide sequence had been identified, a more comprehensive comparison was performed using the β-lactamase database (BLDB) [15]. Figures for the comparison on the genomic context of the β-lactamase genes were produced using clinker [44].

### Phylogenetic analysis

The DNA and corresponding protein sequences of the identified β-lactamase genes were compared to known β-lactamase sequences found in CARD and BLDB (**Supplementary Table 2**). Multiple sequences alignments were performed using T-coffee [45], hosted on the EMBL-EBI webserver [46], to generate both alignments and phylogenetic trees. The phylogenetic trees were visualised using interactive tree of life (iTOL) v3 [47].

### Structural Modelling and Comparison

AlphaFold3 was used for the structural modelling of the predicted β-lactamases [48], and output 5 predicted structures per input protein sequence. The best predicted structures were used for downstream comparisons. Structural comparisons were performed using Matchmaker within ChimeraX, using the Needleman-Wunsch sequence alignment algorithm and the BLOSUM-62 matrix [49]. The structural comparison figures were produced using ChimeraX [50,51].

### β-lactamase activity assay

Overnight cultures of *Pseudomonas* spp. strains were diluted to OD_600_ = 1.0 in BHI broth and transferred to a 96-well plate (190 µL per well). Nitrocefin (1 mg, APExBIO) was resuspended in 0.1 M phosphate buffer pH 7.0 containing 10% DMSO (1 mL). The diluted nitrocefin was added to every well (10 µL) directly prior to assay measurements. The absorbance of each well, at 486 nm, was measured every 90 seconds for 1 hour using a CLARIOstar Plus Microplate reader (BMG Labtech). Negative and positive controls were included on each assay plate, negative controls were nitrocefin in buffer only, and nitrocefin plus diluted *E. coli* NCTC86 culture, while the positive control was nitrocefin with diluted *P. aeruginosa* VS1082 culture. A student’s T-test comparing endpoint absorbance at 1-hour was used, assuming a two-tailed distribution and two-sample equal variance.

### Gibson Cloning

Genomic DNA was extracted from the three *Pseudomonas* isolates using the Wizard^®^ Genomic DNA purification kit (Promega), following the supplied protocol. The resistance genes were amplified by PCR from the extracted DNA, using Q5 DNA polymerase (NEB) for 30 amplification cycles in 25 µL reactions. Cycling conditions were denaturing for 10 seconds (95 °C), annealing for 15 seconds, and extension for 30-seconds (72 °C), except *bla_PFM5_* was a 20-second extension time. The primers were designed to add 20 bp homologous regions of the pEB1 vector during the PCR amplification. Primers and their annealing temperatures used for cloning of resistance genes into pEB1 for heterologous expression are reported in **Supplementary Table S3**, the pEB1 amplification and sequencing primers were taken from [52]. PCR amplification of the pEB1 vector backbone was also undertaken, excluding the sfGFP gene, using 30 cycles in a 50 µL reaction with a 105-second extension time. The NEBuilder^®^ HiFi DNA Assembly Cloning Kit (NEB) was then used to assemble the two fragments together, using a 1:2, vector:insert molar ratio. The assembled plasmids were then transformed into chemically competent NEB5α *E. coli* and selected on kanamycin (30 µg/mL) BHI agar. Colonies containing the correctly assembled plasmids were confirmed by colony PCR and Sanger sequencing, using the pEB1_seq_f primer (**Supplementary Table S3**).

### Minimum Inhibitory Concentration (MIC)

Antibiotics and beta-lactamase inhibitors were purchased from the following suppliers: Apollo Scientific (imipenem), Cayman Chemicals (tazobactam, vaborbactam), Sigma Aldrich (piperacillin), MedChemExpress (ceftazidime, relebactam, xeruborbactam) and ThermoFisher (captopril). MIC values were determined following ISO 20776-1 (2019) guidance using broth microdilution and EUCAST reading guide for broth microdilution version 4.0. Overnight bacterial cultures, grown in BHI, were diluted and OD_600_-corrected to between 0.8-1.0 in cation-adjusted Mueller-Hinton broth (CA-MHB, Sigma Aldrich). They were then further diluted 1:1000 in CA-MHB, which was confirmed to be equivalent to ∼5×10^5^ CFU/mL. Antibiotics for drug testing were serially diluted 1:2 into CA-MHB (final volume 50 µL), in a round-bottom 96-well plate. Diluted bacterial cultures were added to each well (50 µL) and the plates incubated at room temperature (strains did not grow at the standard 37 °C protocol incubation temperature) for 20 hours. Wells were visually inspected for growth, to determine the MIC. MICs were performed in triplicate across three biological replicates, or four biological replicates if there was a discrepancy in the concentrations identified.

## Results

### Screening identifies β-lactam resistant *Pseudomonas* spp

We screened 884 bacterial isolates for growth on both extended spectrum β-lactam (ESBL) supplemented agar and meropenem supplemented agar, and identified seven *Pseudomonas* spp. with presumed β-lactam resistance. We chose to screen against 10 μg/mL meropenem and select for growth on both supplemented agars to investigate isolates with broad β-lactam resistance. Three non-aeruginosa *Pseudomonas* strains (*P. piscis*, *P. saponiphila*, *P. palleroniana*) were selected for follow-up work (**Table 1**), as they represented understudied *Pseudomonas* strains. Their susceptibility was determined against piperacillin, ceftazidime, meropenem and imipenem (**Table 1**) using broth microdilution. All strains were classed as meropenem resistant, with some strains being resistant to all three classes of β-lactams tested, based on the EUCAST MIC breakpoints for non-aeruginosa *Pseudomonas* of 16, 8, 8 and 4 µg/mL of piperacillin, ceftazidime, meropenem and imipenem, respectively.

We performed a phylogenetic comparison of the *Pseudomonas gyrB* gene in our strains against several *Pseudomonas* reference or type strains (**Figure 1**). The *gyrB* gene has previously been used to define novel *Pseudomonas* species or sub-species groups [41,53], and the isolates’ 16S rRNA sequences failed to provide a close match (>98% identity) when run through SpeciesFinder [54]. The phylogenetic tree indicated C001-2B and C001-7H were within the *P. protegens* species group and C006-8D was within the *P. fluorescens* group. Based on average nucleotide identity (ANI) to the genomes of type strains, the closest matches were *P. piscis* for C001-2B, *P. saponiphila* for C001-7H and *P. palleroniana* for C006-8D. MALDI-ToF was also performed on these samples/strains to provide phenotypic analysis of their species. This resulted in a high confidence matches for the C006-8D strain to *P. extremorientalis* and low confidence matches to other *Pseudomonas* species for the other two strains (**Table 1**).

**Figure 1:**
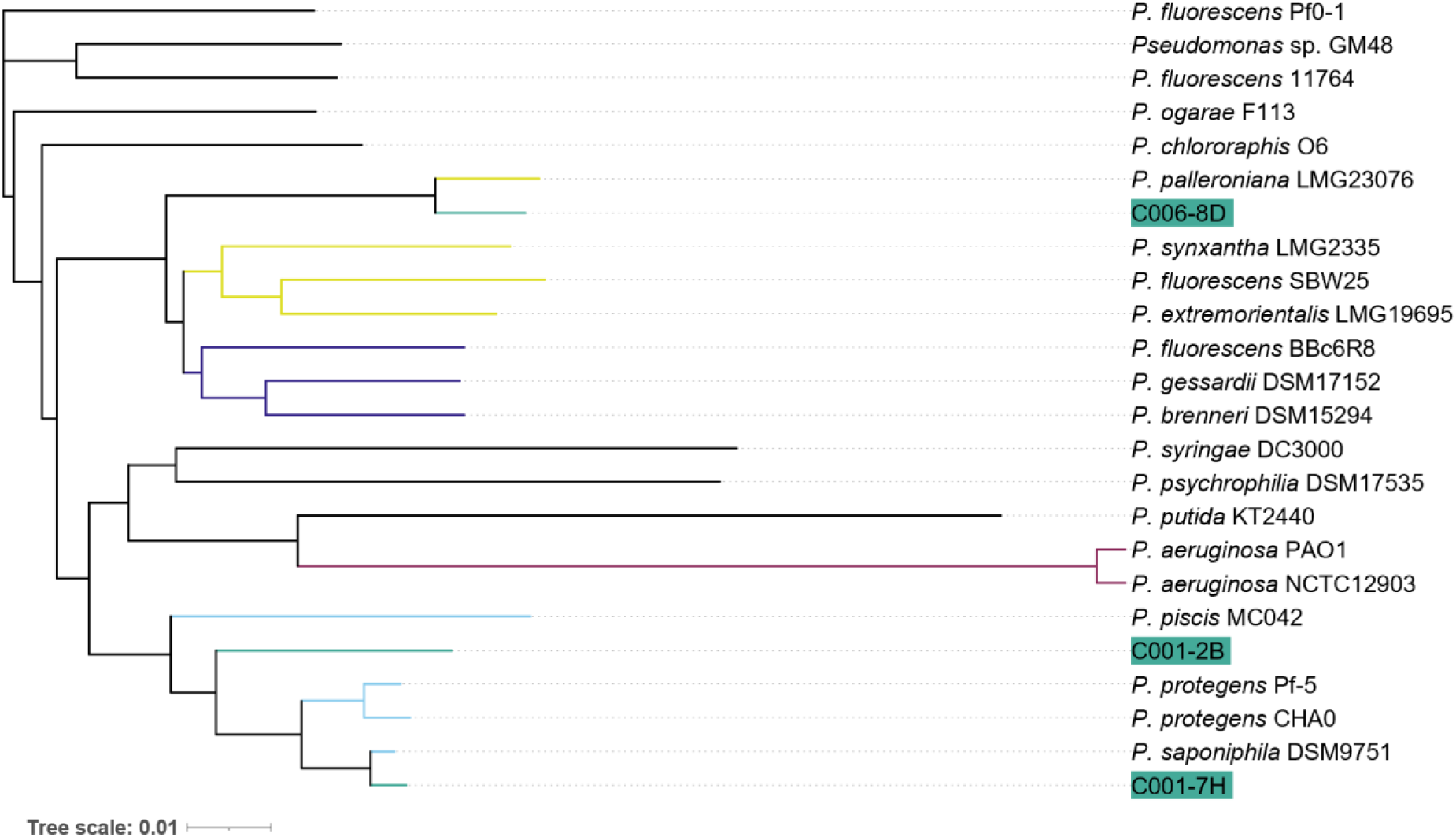
Phylogenetic tree based on the *gyrB* gene sequences of the isolated strains compared with representative *Pseudomonas* spp. The newly isolated strains are highlighted in teal. If multiple *Pseudomonas* strains were from the same overall species group, their branches have been coloured, olive for *P. fluorescens*, indigo for *P. gessardii,* magenta for *P. aeruginosa,* and light blue for *P. protegens*. This phylogenetic tree was generated using T-Coffee and iTOL [45,47].

### Partial database matches to known *bla* genes

We queried the assembled genomes of the three strains on the ResFinder and CARD webservers [12,14], to look for the presence of known ARGs that could confer carbapenem/β-lactam resistance. No β-lactamase genes were identified using the default search settings; however, several non-β-lactamase resistance genes were identified (**Supplementary Table S4**). We then queried the ARG-ANNOT database and two genes, present in C001-2B and C001-7H were found, with between 72-74% nucleotide identity to *bla_PDC-49_* and *bla_PDC-313_*. Further, we detected a sequence in C006-8D with 96% identity to a 55-nucleotide segment of the 765-nucleotide *bla_cphA8_* gene, a subclass B2 metallo-β-lactamase found in *Aeromonas* species.

Prokka annotated both *bla_PDC_*-type genes as ampC [40], and searching the C006-8D genome for ‘ampC’ genes, resulted in the detection an additional putative gene. To increase confidence in these hits, the individual genes were searched in the specialised BLDB database [15], which contained more *Pseudomonas* specific β-lactamase genes. We found matches of 73-76% nucleotide identity to other class C β-lactamase genes. For the subclass B2 nucleotide sequence, this showed a stronger match to the *bla_PFM_* sequences, rather than the short but high identity match to *bla_cphA8_*.

### Novel β-lactamase sequences show similarity to known genes

To investigate where these novel sequences fit into the broader β-lactamase landscape, we compared the DNA sequences within each class or subclass of β-lactamase (**Figure 2**). Representatives of all known sub-classes of class C β-lactamase genes were compared with the three novel genes we identified (**Figure 2a**). While the *bla_PDC_* genes were closely related to the novel genes, *bla_PFL-1_* was found to be more closely related, especially to the novel gene in C006-8D. The latter identified in the environmental *P. yamanorum*, compared to all the former genes found in *P. aeruginosa* strains. We have named these three novel genes as *bla_PFL7_*, *bla_PFL8_* and *bla_PFL9_*, based on all three strains being part of the *P. fluorescens* complex and *bla_PFL1-6_* already being assigned in BLDB. In contrast to the diversity of class C β-lactamases, the subclass B2 metallo-β-lactamases have only five distinct groups and only 24 currently reported sequences. Comparison of the C006-8D-B2 sequence indicates it has the highest identity to *bla_PFM-1_* (**Figure 2b**), at 87.8% nucleotide identity. Thus, this gene has been assigned as *bla_PFM-5_* by NCBI (PV244044.1).

**Figure 2:**
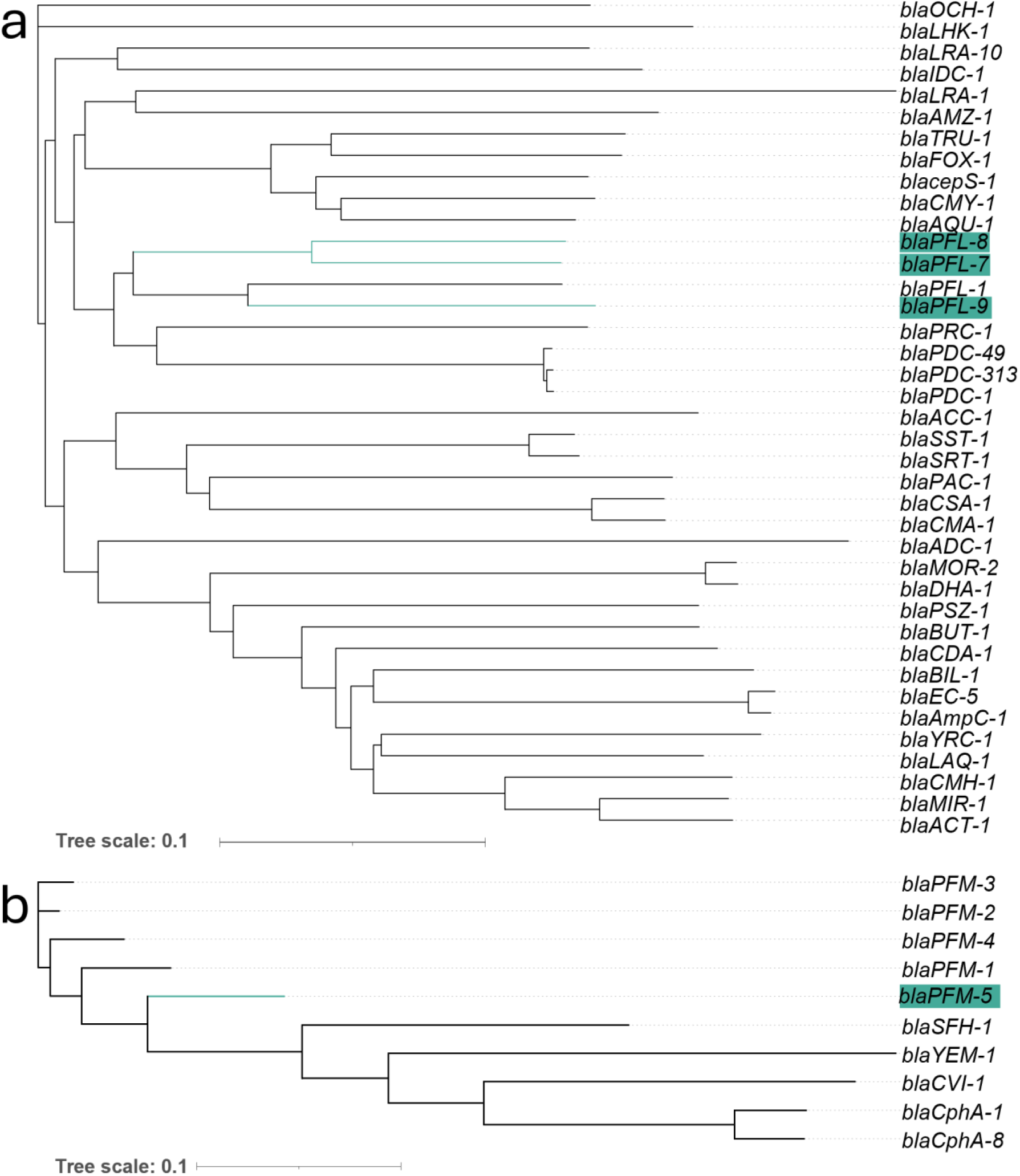
Comparison of nucleotide sequences encoding (a) class C β-lactamases and (b) subclass B2 β-lactamases. Highlighted in teal are the novel β-lactamase genes found within this work.

To evaluate potential functionality of the β-lactamase DNA sequences, we compared the translated protein sequences with previously characterised or identified β-lactamase proteins. A phylogenetic tree was created, with representatives of all class C β-lactamases in the CARD database [12] (**Supplementary Figure S1**). From this, a subset of class C β-lactamases was selected for multiple sequence alignment with the novel β-lactamases reported herein (**Supplementary Figure S2a**). The five residues required for catalytic activity (Ser64, Lys67, Tyr150, Asn152 and Lys315) in class C β-lactamases were retained in our three novel sequences [33,55]. The sequences directly surrounding these catalytic residues were also highly conserved. The closest matches to these novel genes were all from environmental *Pseudomonas* strains.

We also created a phylogenetic tree comparing the protein sequences of the subclass B2 β-lactamases, indicating the closest match was BlaPFM-1 at 87.8% identity, which was found in *P. fluorescens* [29]. (**Supplementary Figure S1b**). Multiple sequence alignment was also performed to compare the novel subclass B2 metallo-β-lactamase with the other reported sequences (**Supplementary Figure S2b**). Six residues (Asn116, His118, Asp120, His196, Cys221, His263) are conserved across all subclass B2 β-lactamase, being required for zinc cofactor binding or catalytic activity [30]. These six residues, among many other residues, were conserved in the novel β-lactamase we identified.

### AlphaFold structural predictions are highly similar to comparable crystal structures

We predicted the structures of all three class C β-lactamases and the subclass B2 β-lactamase using AlphaFold3 [48], with the majority of each protein having a very high confidence prediction (>90, **Supplementary Figure S3**). The one region with a low AlphaFold confidence score (<50) was the N-terminal tail region, representing a weak structural prediction. Comparing the best ranked model to each other indicated a high level of structural similarity, despite the lower sequence similarity (**Figure 3a**). The root-mean-square-deviation (RMSD) between BlaPFL7 and BlaPFL8 was 0.417 Å across the pruned alpha-carbon atoms of the proteins, and between BlaPFL7 and BlaPFL9 was 0.461 Å. These RMSD values are likely to be elevated due to the N-terminal region being included in the calculations.

**Figure 3:**
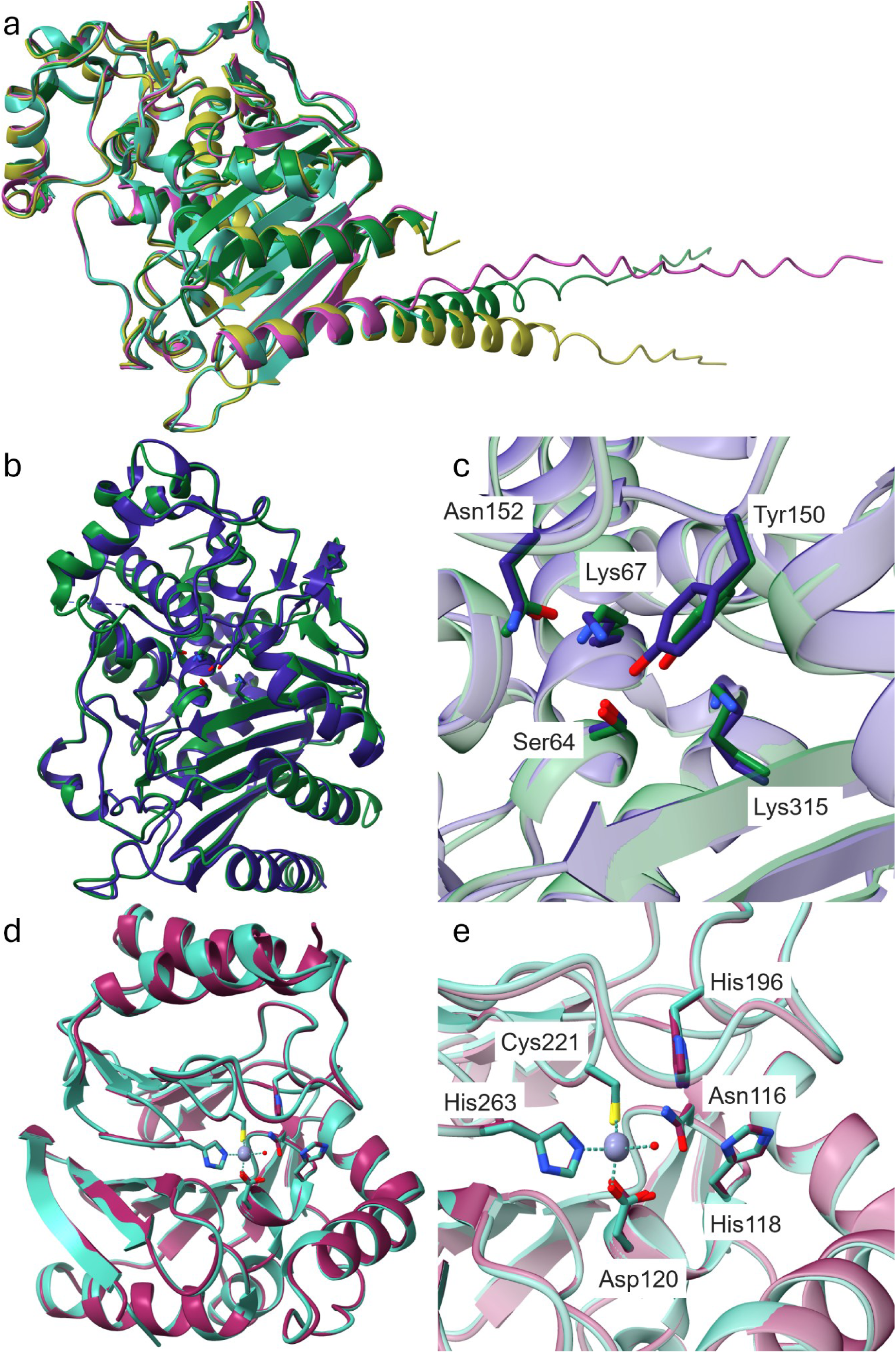
AlphaFold structural predictions of the three novel class C and one novel subclass B2 β-lactamases show strong similarity to each other, and to the previously solved AmpC-type/subclass B2 crystal structures (2QZ6 and 3SD9). (a) Overlay of the three AlphaFold models, showing the best prediction for each class C protein sequence (b) Comparison of BlaPFL7 (green) to 2QZ6 (indigo), excluding BlaPFL7 N-terminal amino acids 1-27 for clarity. (c) Comparison of BlaPFL7 to 2QZ6 active site, highlighting the similar 3D-orientation of the catalytic residues (Ser64, Lys67, Tyr150, Asn152 and Lys315). (d) Comparison of the BlaPFM5 modelled structure (magenta, terminal amino acids 1-23 removed for clarity) to the crystal structure of SFH-1 (cyan). (e) Comparison of BlaPFM5 to 3SD9 active sites, highlighting similar 3D-orinetation of subclass B2 conserved residues (Asn116, His118, Asp120, His196, Cys221, His263 [30]). Figure was produced using ChimeraX.

The predicted structures were also compared to the nearest homologue with a crystal structure, for class C proteins this was BlaPFL-P1 (2QZ6) [33]. For BlaPFL7, the RMSD was 0.603 Å (across 345 atom pairs), indicating that the models are similar to the crystal structure (**Figure 3b**). The structural comparisons of the other two modelled proteins to BlaPFL-P1 are displayed in **Supplementary Figure S4**. In addition, the catalytic residues were predicted to have similar 3D-orientations to the BlaPFL-P1 crystal structure (**Figure 3c**). For the metallo-β-lactamase gene, we compared the modelled protein structure to SFH-1 (3SD9), the closest homologue match to have been crystallised. The RMSD to the SFH-1 crystal structure was 0.479 Å across all 229 alpha carbon pairs (**Figure 3d**). The six catalytic residues required for catalytic activity and zinc ion coordination were predicted to retain similar 3D orientation in the active site (**Figure 3e**). In addition, AlphaFold3 also predicted the same zinc coordination displayed in the SFH-1 crystal structure.

### All three *Pseudomonas* spp. strains display β-lactamase activity

Given the high predicted structural similarity and conservation of required catalytic residues, we assessed the catalytic ability of the β-lactamases to degrade the cephalosporin nitrocefin. The degradation of nitrocefin results in a shift in absorbance from 390 nm to 486 nm, alongside a colour change from yellow to red. To control for the breakdown of nitrocefin within BHI, a negative control of *E. coli* NCTC86 was utilised, as it possessed no β-lactamases genes. As a positive control, the *P. aeruginosa* VS1082 strain was used, encoding both a *bla_PDC_* and a *bla_OXA486_* β-lactamase. All three strains could degrade nitrocefin within 1-hour, p<0.001 comparing assay end-point absorbance to *E. coli* control (**Figure 4**). Strain C001-7H had the highest activity, causing an increase in absorbance at 486 nm of 0.800±0.113, compared to an increase of 1.147±0.230 for *P. aeruginosa* VS1082 and 0.023±0.043 for the *E. coli* negative control. In contrast, the least active strain, C001-2B, caused an increase in absorbance of 0.385±0.065. The highest activity of strain C001-7H against nitrocefin also correlated with the highest MICs against the cephalosporin ceftazidime.

**Figure 4:**
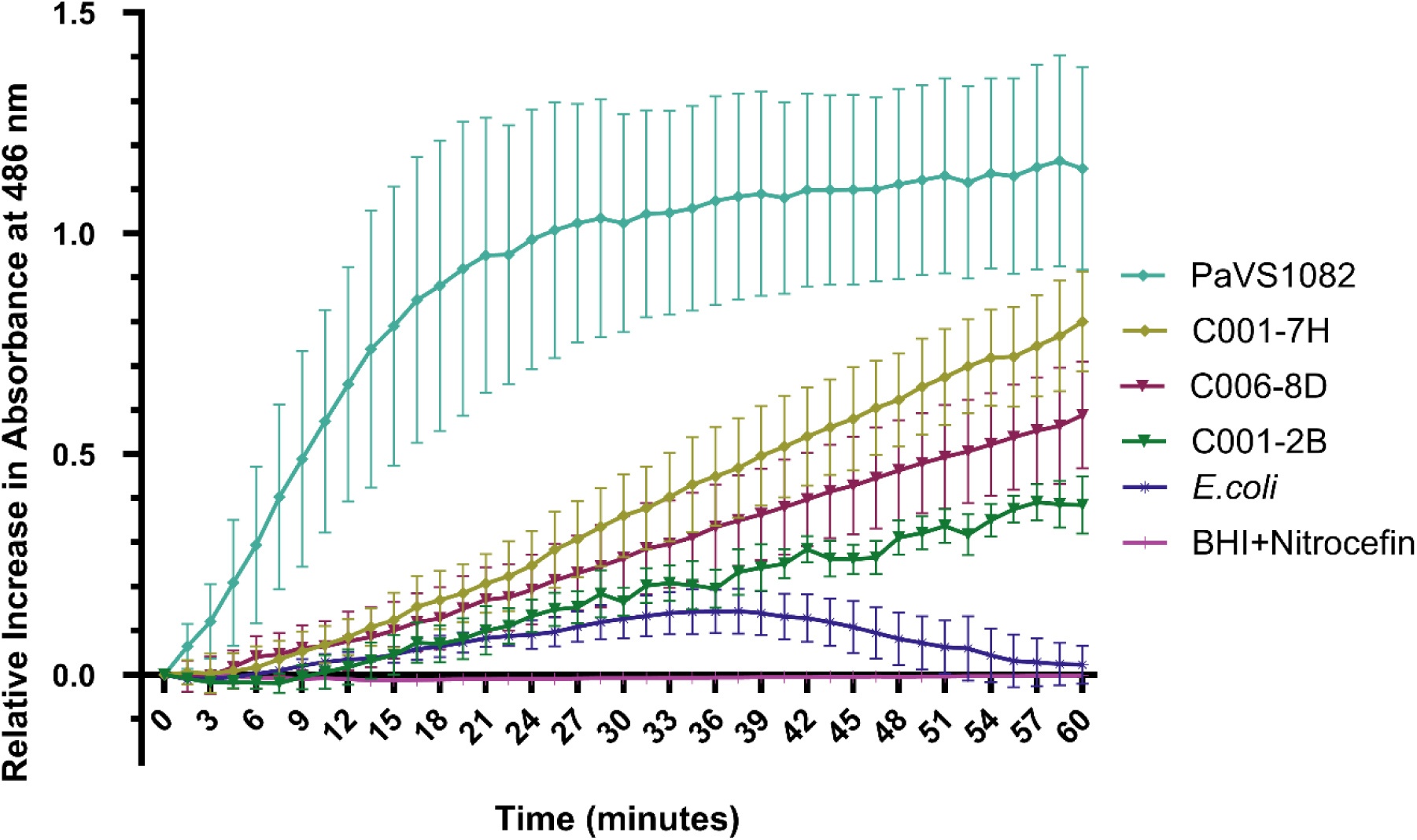
Assessing the resistant mechanisms of β-lactam resistant strains. (A) Nitrocefin activity assay showing all three Pseudomonas strains have the ability to degrade nitrocefin. *P. aeruginosa* VS1082 (PaVS1082) is used as positive control with known β-lactamases (*bla_PDC_* and *bla_OXA486_*), while the *E. coli* NCTC86 is used a non-β-lactamases producing negative control. Three biological repeats, each with three technical replicates.

### Ectopic expression of novel *bla* genes confer resistance in *E. coli*

Given the ability of these β-lactamases to cleave nitrocefin and the strains being resistant to a broad range of β-lactams, we cloned these novel genes into an *E. coli* NEB5α strain to test these gene in a clean and different genetic backgound. Each gene was individually cloned into the low copy number expression vector pEB1-*sfGFP* [56]. The genes were put under constitutive expression of the *proC* promoter, by replacing the *sfGFP* with the β-lactamase genes. We then tested antibiotic susceptibilities of the pEB1-*bla* transformed *E. coli* and the susceptible *E. coli* NEB5α laboratory strain using broth microdilution for piperacillin, ceftazidime, meropenem and imipenem. In addition, we tested several β-lactamase inhibitors: tazobactam, relebactam, vaborbactam, xeruborbactam and captopril.

*E. coli* NEB5α strains containing the pEB1 plasmids encoding the novel β-lactamases could confer resistance to either piperacillin and ceftazidime, or to carbapenems, but not to all three groups (**Table 2)**. *bla_PFL7_*, *bla_PFL8_* and *bla_PFL9_* conferred resistance above EUCAST’s clinical breakpoints to piperacillin and ceftazidime. However, these three genes did not confer resistance to either carbapenem tested. The addition of tazobactam was able to restore susceptibility to piperacillin for all three genes. In contrast, *bla_PFM5_* conferred imipenem resistance and intermediate resistance to meropenem, confirming carbapenemase activity. This gene lacked the ability to confer piperacillin or ceftazidime resistance. The addition of relebactam, vaborbactam, xeruborbactam or captopril, as known metallo-β-lactamase inhibitors, had no impact on the carbapenem resistance of the *E. coli* strain.

### High amino acid identity matches are present in publicly available sequencing data

We queried the translated amino acid sequences of *bla_PFL_* and *bla_PFM_* genes for similar proteins with >95% amino acid identity using both the non-redundant protein sequences and metagenomic proteins databases with BLASTp [43]. BlaPFL7 had two matches at 100% identity (100% coverage, WP_226524014.1), with no other matches >95% identity. The identical protein sequences were present in *Pseudomonas* spp., isolated in 2021 from air in a pyrite mine in Lousal, Portugal and from a metagenomic assembled genome derived from a cow manure metagenome from the Czech Republic collected in 2018 [57,58]. BlaPFL8 had six matches with between 98.7% and 99.2% identity (100% coverage, WP_265533797.1, WP_110724755.1 and WP_092312625.1). All similar protein sequences were present in *P. saponiphila* or unclassified *Pseudomonas* spp., the former found in Georgia and Michigan, USA, while the latter were all found in pond water from University of Southern Alabama campus, USA. BlaPFL9 had a match with 94.7% identity (100% coverage, WP_090370723.1), in a *P. palleroniana* isolate from Cameroon in 1998 from diseased rice seeds and leaf sheaths. Finally, BlaPFM5 had 18 matches at 94.47% identity (100% coverage, WP_102591908.1), with no other matches >91% identity. All these matches were from related isolates found in groundwater in Oak Ridge, Tennessee, USA [59]. The metagenomic proteins database only returned low identity matches with >90% coverage; the highest percentage identities of 64% for BlaPFL7, 62% for BlaPFL8, 79% for BlaPFL9 and 56% for BlaPFM5, from either compost or marine metagenomes. Overall, this suggests these, and variants of these genes are geographically widespread amongst environmental *Pseudomonas* spp., but there were very low numbers of matches in general. As there is a database bias towards clinically relevant pathogens and their associated ARGs, this highlights environmental settings as understudied reservoirs of antimicrobial resistance.

### β-lactamase genes are found in different genomic contexts

We looked at the genomic context of these genes, to explore if the novel β-lactamase genes are associated with mobile genetic elements (MGEs) with the potential to move from environmental isolates into potentially pathogenic *Pseudomonas*. One approach was to look at the %GC content of the gene relative to the genome average, which could suggest recent movement into the strain’s genome [30]. The class C β-lactamase genes had similar %GC content to the genomes, *bla_PFL7_* had 64.4% compared to 63.3% of the C001-2B genome, *bla_PFL8_* had 64.6% compared to 61.9% of the C001-7H genome and *bla_PFL9_* had 60.1% compared to 59.9% of the C006-8D genome. The exception was *bla_PFM5_* which had 53.4% GC compared 59.9% in the genome and hence was investigated further.

This was mirrored by the wider 33.8 Kb contig that the novel *bla_PFM5_* gene was within, having an overall 53.6% GC content. MobileElementFinder was used to search for the presence of MGEs on this contig [60], but resulted in no significant matches. A low-quality match to IS*30* was displayed when including inferred transposons, showing matches at 96.7% identity but only 11.4% coverage at both ends of the contig. Searching using ISFinder [61], both nucleotide sequences were found to match to IS*Ppu17* (IS*30* family) with 97% identity across the first 122 nucleotides of the 1,066 nucleotide IS element, until the end of the contig (**Supplementary Figure S5**). The putative IS element at the 5’ end of the contig was directly preceded by a truncated *ompP1*/*fadL* transporter gene (**Supplementary Figure S5**). The genome was also analysed by IslandViewer4, to search for genomic islands [62]. This identified two putative genomic islands using SIGI-HMM of 4,249 bp and 5,031 bp on the same contig as the *bla_PFM5_* gene, the closest being 7,048 nucleotides upstream.

Finally, we explored the specific genomic context surrounding this novel gene, relative to the previously reported *bla_PFM1_* and *bla_PFM2_* and *bla_PFM4_* genes (**Figure 5a**). The sequence of *bla_PFM3_* was excluded, as only the nucleotide sequence of the specific gene was published [29]. There are many high-percentage identity matches between genes surrounding the subclass B2-metallo-β-lactamase genes. Many of the conserved genes upstream of *bla_PFM5_* were predicted to encode type six secretion system (T6SS) components, based on gene annotation, including *tssA*, *tssB*, *tssC*, *hcp1*, *tssE*, *tssF*, *tssG*, *vgrG*, *tssJ*, *tssK*, *tssL* and *tssM* [63,64]. The only required T6SS gene not present in these gene clusters is *clpV*, although two copies are found in strain C006-8D, elsewhere in the genome.

**Figure 5:**
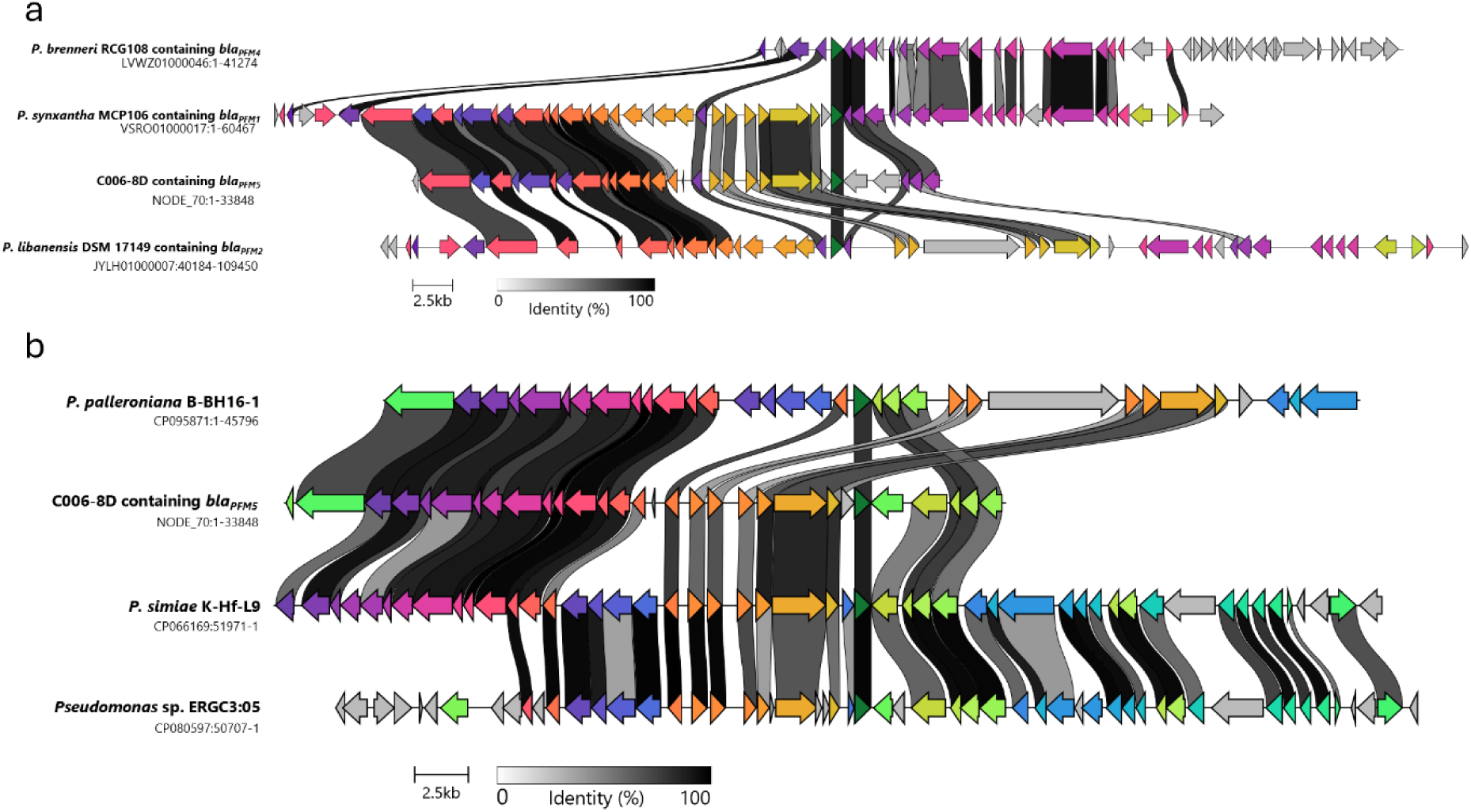
(a) Genomic context of *bla_PFM5_* in comparison to the three other *bla_PFM_* genes, displaying the whole contigs for *bla_PFM4_* and *bla_PFM5_*, and segments of *bla_PFM1_* and *bla_PFM2_*. (b) Genomic comparison of the contig containing *bla_PFM5_* to segments of other *Pseudomonas* spp. genomes identified through BLASTn searching for homologous genes. In (a) and (b), the *bla_PFM_* genes are highlighted in dark green and centralised, the other genes are coloured if there was >30% identity between two or more of the input genes across separate contigs.

As there was variation in the genes surrounding *bla_PFM5_* when compared to the other *bla_PFM_* genes, we investigated whether conserved sections of the contig were present in other WGS data. We used BLASTn to search for matches to the group of seven genes directly preceding the *bla_PFM5_* gene, as they were found as a conserved group in the C006-8D and *P. synxantha* MCP106 genomes [43]. This resulted in three hits, all within environmental *Pseudomonas* spp. genomes [65–67]. All three of these hits also contained a *bla_PFM_* type gene, downstream of the matching query genes. These putative subclass B2-metallo-β-lactamase genes had the highest nucleotide identity to the *bla_PFM2_* gene, with between 87.5 to 97.8%. These genomic regions also showed clear differences in gene content and arrangement compared to the contig we found containing the *bla_PFM5_* gene (**Figure 5b**) which could suggest genetic divergence over time.

## Discussion

This study has identified four novel, experimentally confirmed, β-lactamase genes which are present in non-aeruginosa *Pseudomonas* spp. found in untreated wastewater. Three of these genes were class C β-lactamases, *bla_PFL7_*, *bla_PFL8_*, and *bla_PFL9_*, while the fourth gene was a sub-class B2 metallo-β-lactamase, *bla_PFM5_*. These genes were not readily identifiable when screening against resistance gene databases, which typically focus on resistance genes found in clinically relevant bacterial hosts.

The previously characterised *bla_PRC1_* and *bla_PFL-P1_* genes were the most similar to the novel class C β-lactamases reported herein. A direct comparison to *bla_PRC1_* is possible due to its previous cloning into *E. coli* DH5α, where it was unable to confer resistance to either piperacillin or ceftazidime [35], unlike the *bla_PFL7-9_* genes we report here. However, this gene was able to confer resistance to other penicillins and cephalosporins, including 8-fold increases in MIC for ampicillin and cefotaxime, indicating a different spectrum of activity [35]. BlaPRC-1 was also unaffected by β-lactamase inhibitor tazobactam, in contrast to all three of the reported BlaPFL proteins being inhibited. For BlaPFL-P1, only kinetic parameters for the purified enzyme were determined, preventing direct activity comparisons on a cellular level [33].

Comparing the *bla_PFM5_* β-lactamase to the closest matches *bla*_PFM-1-4_, they all show specific carbapenemase activity but not penicillinase or cephalosporinase activity [29,30], a defining feature of subclass B2. When previously cloned into susceptible *E. coli*, *bla*_PFM-1-4_ genes showed increased resistance to carbapenems; imipenem and meropenem (**Table 3)** [29,30], similarly to our described *bla_PFM5._* Greater resistance to imipenem relative to meropenem was conferred by b*la*_PFM-2_ and *bla*_PFM-3_, 4-fold and 8-fold higher MIC values, while *bla*_PFM-4_ led to 2-fold higher MIC values against meropenem. Despite higher sequence identity to *bla*_PFM-1_, the *bla_PFM-5_* encoded carbapenemase conferred a 2-fold higher MIC to imipenem over meropenem. BlaPFM-5 was also unaffected by all clinically used metallo-β-lactamase inhibitors, and captopril, which is being investigated for this application [68–71]. Both xeruborbactam and captopril have shown inhibitory activity against clinically relevant subclass B1 metallo-β-lactamases [68,69], but do not inhibit the subclass B2 metallo-β-lactamase BlaPFM-5 reported here.

**Table 3:** Carbapenem resistance conferred by *bla_PFM_* genes when cloned into susceptible *E. coli*.

| Gene | Meropenem MIC<br>(μg/mL) | Imipenem MIC<br>(μg/mL) | Data source |
| --- | --- | --- | --- |
| *bla*<sub>PFM-1</sub> | 1 | 1 | Poirel et al. 2020 [29] |
| *bla*<sub>PFM-2</sub> | 1 | 4 | Poirel et al. 2020 [29] |
| *bla*<sub>PFM-3</sub> | 2 | 16 | Poirel et al. 2020 [29] |
| *bla*<sub>PFM-4</sub> | 32 | 16 | Cafiero et al. 2022 [30] |
| *bla*<sub>PFM-5</sub> | 4 | 8 | This work |

The cloned class C β-lactamases were unable to confer carbapenem resistance within *E. coli*, however, the isolated *Pseudomonas* spp. strains were meropenem resistant. This suggests the strains may have other resistance mechanisms, such as efflux or decreased uptake. Both resistance mechanisms have previously been implicated in β-lactam resistance in *Pseudomonas*, such as MexAB-OprM efflux pump or mutations in the OprD membrane porin which reduce imipenem entry [16,72,73]. However, drug-resistant efflux pumps were not identified by CARD or ResFinder in these three *Pseudomonas* spp. (**Supplementary Table S4**), which could be due to lack of presence or lack of sequence homology. The resistance to imipenem in C001-2B could indicate OprD mutations in this strain. In the isolated *Pseudomonas* spp., there are multiple copies of the *oprD* genes, five for each of C001-2B and C001-7H and eight in C006-8D. The encoded proteins had amino acid sequence identities of between 31.4% to 60.9% to the *P. aeruginosa* PA01 (ATCC 15692) OprD sequence (P32722) [74].

In relation to the mobility of these novel β-lactamases, the use of short-read sequencing has limited our ability to confirm the presence of MGEs. All three of the class C β-lactamases appear to be in the middle of large contigs, with no known MGEs nearby, suggesting that they have low potential for mobility within their chromosomal locations. As the genes are from environmental *Pseudomonas* spp., an IS element could be present, but which has not been characterised or included in the searched databases. For the *bla_PFM5_* gene, the smaller contig size and lower GC content than the rest of the genome may suggest a transposition event in the past. In addition, the truncated *ompP1*/*fadL* gene directly preceding the putative IS element on this contig may have occurred due to an insertion at this site. However, due to the lack of a full length IS element sequence present near the *bla_PFM5_* gene, we cannot prove the presence of a MGE within the current work.

Comparing the genomic context directly surrounding all the *bla_PFM_* genes, there were clear differences in the number, location and conservation of genes. As the *bla_PFM_* genes were from a range of *Pseudomonas* spp., this could represent divergence over time or transposition occurring within genomes or between species. There were also gaps in the nucleotide alignment between our contig and those identified through BLAST, in the regions directly preceding and following the *bla_PFM_* genes. Our ability to identify additional putative subclass B2 metallo-β-lactamase genes, using adjacent genes found on the same contig as the *bla_PFM5_* gene, provides additional evidence of their widespread distribution within environmental *Pseudomonas* spp. Associated with the *bla_PFM_* genes, we found almost complete sets of T6SS genes, excluding *clpV*, in three of the other *Pseudomonas* spp. strains and partial sets in two *Pseudomonas* spp. isolates. T6SSs have been associated with virulence and interbacterial competition [64].

Our study highlights our limited knowledge of environmental ARGs, especially when they have lower sequence identity to clinically relevant ARGs present within public databases. Without the associated phenotypic resistance data, it is unlikely that genes with nucleotide sequence identity as low as 70% to known ARGs would be investigated. Hence, metagenomic analysis of environmental communities may underestimate the prevalence of ARGs and ignore novel sequences with potential future clinical relevance. This emphasises the importance of experimental validation of putative ARGs and our study outlines an approach for studying novel β-lactamase genes.

## Author Statements

### Authors and Contributors

AK and APR; conceptualisation. AK, EA, AB, KD, CM, and FG; investigation. AK, EA and FG; methodology. AK; data curation. AK, AB, KD and FG; resources. AK; writing – original draft. All authors; writing – review & editing. APR; supervision. APR; funding acquisition.

## Acknowledgments

pEB1-sfGFP plasmid was a kind gift from Philippe Cluzel (Addgene: http://n2t.net/addgene:103983). *E. coli* NCTC86 was a gift from the National Collection of Type Cultures

## Data Availability

The *bla_PFM-5_* nucleotide sequence is available at PV244044.1, with the translated amino acid sequence at XPT09377.1. The whole genome sequencing data is available as part of BioProject PRJNA1161700. The raw reads have been deposited in SRA under the accession numbers SRR36506414, SRR36506413 and SRR36506412. The assembled and annotated genome has been deposited in GenBank under accession numbers JBTANL000000000, JBTANK000000000 and JBTANJ000000000.

