## Supplementary Tables and Figures for "Novel Class B2 and C β-lactamases harboured by *Pseudomonas* spp. wastewater isolates"

### Supplementary Material

**Supplementary Table S1:** Accession numbers of *gyrB* nucleotide sequences and associated accession numbers

| Species | Genome accession number | *gyrB* accession number |
|---|---|---|
| *P. aeruginosa* NCTC12903 | LR134309.1 | AAG03394.1 |
| *P. aeruginosa* PAO1 | AE004091.2 | AAG03394.1 |
| *P. brenneri* DSM15294 | VFIL01000014.1 | TWR75120.1 |
| *P. chlororaphis* O6 | AHOT01000001.1 | EIM18870.1 |
| *P. extremorientalis* LMG19695 | NZ_MDGK01000047.1 | WP_025854386.1 |
| *P. fluorescens* 11764 | CP010945.1 | AKV10789.1 |
| *P. fluorescens* BBc6R8 | CP065593.1 | QQD53483.1 |
| *P. fluorescens* Pf0-1 | CP000094.2 | ABA71748.1 |
| *P. fluorescens* SBW25 | OV986001.1 | CAI2794345.1 |
| *P. gessardii* DSM17152 | MNPU01000041.1 | ONH38849.1 |
| *P. ogarae* F113 | CP003150.1 | AEV60038.1 |
| *P. palleroniana* LMG23076 | PYWX01000029.1 | PTC28769.1 |
| *P. piscis* MC042 | WHUV01000001.1 | MQA51704.1 |
| *P. protegens* CHA0 | CP003190.1 | AGL81811.1 |
| *P. protegens* Pf-5 | CP000076.1 | AAY95422.1 |
| *P. psychrophilia* DSM17535 | JYKZ01000013.1 | KMM97129.1 |
| *P. putida* KT2440 | AE015451.2 | AAN65647.1 |
| *P. saponiphila* DSM9751 | FNTJ01000002.1 | SEC75098.1 |
| *P. synxantha* LMG2335 | BDAK01000011.1 | WP_043048232.1 |
| *P. syringae* DC3000 | AE016853.1 | AAO53561.1 |
| *Pseudomonas* sp. GM48 | AKJM01000051.1 | EJM59090.1 |

**Supplementary Table S2:** Accession number of β-lactamase sequences utilized in Figure 2 and Supplementary Figure S1

| B-lactamase gene ID | GenBank ID | GenPept ID |
|---|---|---|
| **AmpC-type β-lactamase genes** | | |
| *blaACC-1* | AJ133121 | CAB46491 |
| *blaACT-1* | U58495 | AAC45086 |
| *blaADC-1* | AY228469 | AAR00515 |
| *blaAmpC-1* | FN649414 | CBJ02047.1 |
| *blaAMZ-1* | CP123364 | WGJ88903 |
| *blaAQU-1* | AB765393 | BAM76828 |
| *blaBIL-1* | X74512 | CAA52618 |
| *blaBUT-1* | AJ415568 | CAC94553 |
| *blaCDA-1* | KJ650399 | AID52933 |
| *blaCepS-1* | X80277 | CAA56561 |
| *blaCMA-1* | KF640251 | AHB24364 |
| *blaCMH-1* | JQ673557 | AFI56422 |
| *blaCMY-1* | X92508 | CAA63264 |
| *blaCSA-1* | KF623543 | AHB24365 |
| *blaDHA-1* | JN638038 | AEP68014 |
| *blaEC-5* | DQ092424 | AAZ85965 |
| *blaFOX-1* | X77455 | CAA54602 |
| *blaIDC-1* | MN985649 | QIB98918 |
| *blaLAQ-1* | MZ497396 | QXM27670 |
| *blaLHK-1* | AY632070 | AAT46340 |
| *blaLRA-1* | EU408346.1 | ACH58980.1 |
| *blaLRA-10* | EU408357 | ACH58999 |
| *blaMIR-1* | M37839 | AAD22636 |
| *blaMOR-2* | AY235804 | AAO84036 |
| *blaOCH-1* | CP008820 | AIK44884 |
| *blaPAC-1* | KY285014 | APM84516 |
| *blaPDC-1* | AY083595 | AAM08945 |
| *blaPDC-313* | MH780090 | AXQ11882 |
| *blaPDC-49* | KJ949064 | AIG19986 |

| | | |
|---|---|---|
| *blaPFL-1* | CP012400 | AMW83344 |
| *blaPRC-1* | MW854031 | QTV22830 |
| *blaPSZ-1* | OQ725878 | WGF22896 |
| *blaSRT-1* | AB008454 | BAA23130 |
| *blaSST-1* | AB008455 | BAA23131 |
| *blaTRU-1* | EU046614 | ABW05394 |
| *blaYRC-1* | DQ185144 | ABA70720 |
| **Subclass B2 β-lactamase genes** | | |
| *blaCphA-1* | X57102 | CAA40386 |
| *blaCphA-8* | AY261375 | AAP97129 |
| *blaCVI-1* | NG_067986 | WP_164461288 |
| *blaPFM-1* | MN065826 | QDC33502 |
| *blaPFM-2* | JYLH01000007 | KRP45299 |
| *blaPFM-3* | MN080497 | QGJ02507 |
| *blaPFM-4* | LVWZ01000046 | OAE14554 |
| *blaSFH-1* | NZ_AUZV01000091 | WP_024531368 |
| *blaYEM-1* | CTIO01000005 | CQH06348 |

**Supplementary Table S3:** Primers used for PCR cloning and sequence confirmation of novel resistance genes into pEB1. Lower case sequences match with the vector, while upper case sequences match with the inserted genes.

| Primer | Purpose | Sequence (5’ -> 3’) | Annealing temperature (°C) |
|---|---|---|---|
| pEB1_seq_f | Sequencing primer - Forward | gcgtatcacgaggccctttc | 52 |
| pEB1_seq_r | Sequencing primer - Reverse | aaagggaaaactgtccatatgcac | 52 |
| pEB1_f | Vector Amplification - Forward | taaatgtccagacctgcagg | 57 |
| pEB1_r | Vector Amplification - Reverse | atgtatatctccttcttaaatctag | 57 |
| C001-2B_f_pEB1 | Insert Amplification - Forward | tttaagaaggagatatacatATGCGGCAAACAACCTTG | 64 |
| C001-2B_r_pEB1 | Insert Amplification - Reverse | cctgcaggtctggacatttaTTACCTGGCGCTGTCCAG | 64 |
| C001-7H_f_pEB1 | Insert Amplification - Forward | tttaagaaggagatatacatATGCGGCATACAACCTTAAC | 62 |
| C001-7H_r_pEB1 | Insert Amplification - Reverse | cctgcaggtctggacatttaTTACTTCGCAGTCTCCAG | 62 |
| C006-8D_AmpC_f_pEB1 | Insert Amplification - Forward | tttaagaaggagatatacatATGAATGCCCCACTCCAAAAATTTC | 67 |
| C006-8D_AmpC_r_pEB1 | Insert Amplification - Reverse | cctgcaggtctggacatttaTCAGTGGTCCATGGCACTC | 67 |
| C006-8D_CphA_f_pEB1 | Insert Amplification - Forward | tttaagaaggagatatacatATGACATTGACTAAACTCCTC | 57 |
| C006-8D_CphA_r_pEB1 | Insert Amplification - Reverse | cctgcaggtctggacatttaTTACTGCTTGTCGTGCTC | 57 |

**Supplementary Table S4:** Non-β-lactamase genes identified by CARD, ResFinder and ARG-ANNOT. All analysis run on default settings, displaying outputs with >60% identity (>90% for ResFinder based on default cut-off). id = identity.

| Strain | CARD output | ResFinder Output | ARG-ANNOT output |
|---|---|---|---|
| C001-2B | yajC – 90% id (100% length)<br>fosA – 72% id (102% length)<br>adeF – 67% id (100% length) | No matches | oqxBgb – 75% id (1361/3153 nt) |
| C001-7H | qacE – 98% id (100% length)<br>yajC – 91% id (100% length)<br>fosA – 72% id (102% length) | qacE – 100% id | No matches |
| C006-8D | abaQ – 73% id (101% length)<br>soxR – 72% id (94% length)<br>adeF – 66% id (81 % length)<br>fosA8 – 59% id (96% length) | No matches | oqxBgb – 72% id (1495/3153 nt) |

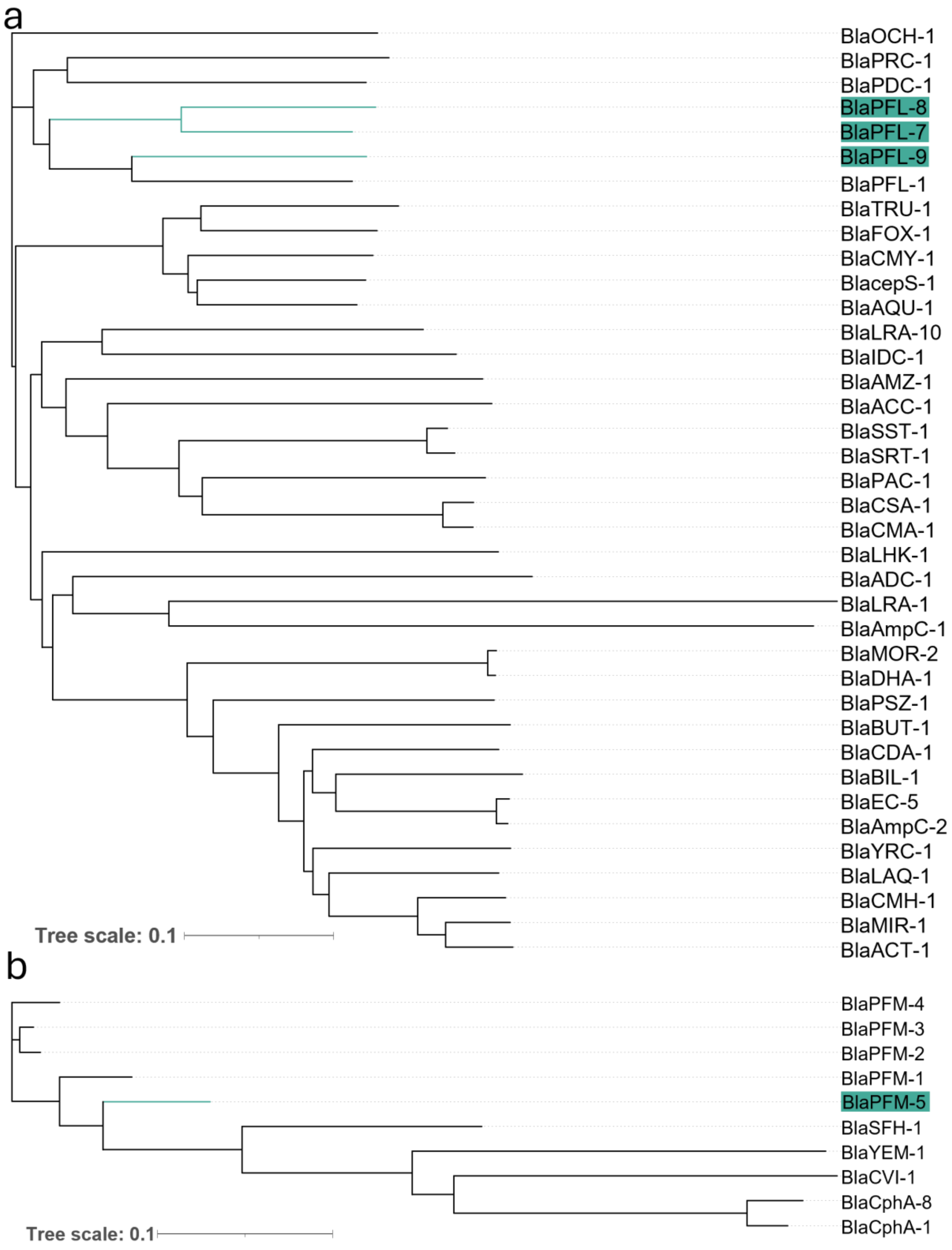


**Supplementary Figure S1**: Phylogenetic trees comparing the (a) class C β-lactamase or (b) subclass B2 metallo-β-lactamase protein sequences. Highlighted in teal are the novel β-lactamases discovered in this work.

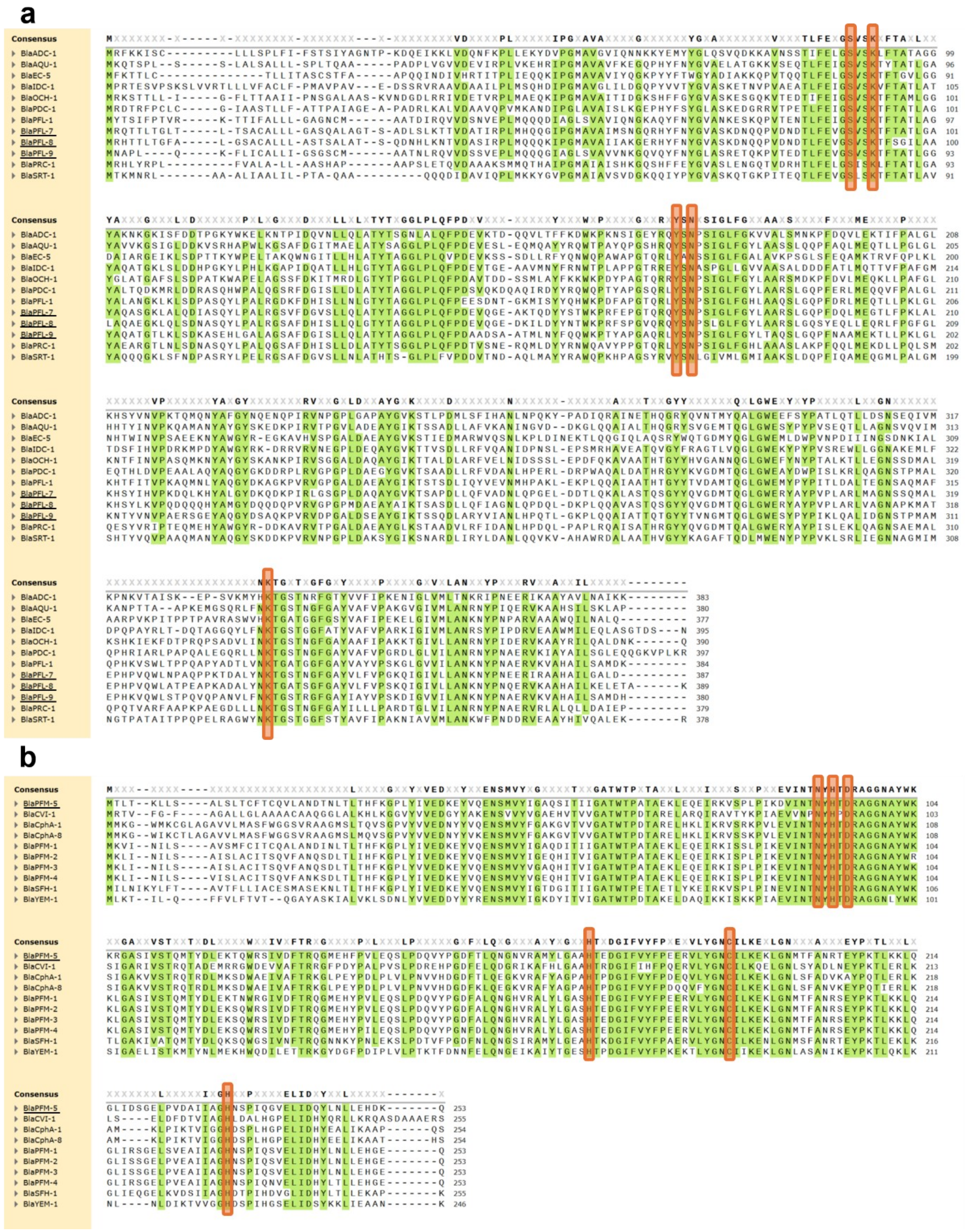


**Supplementary Figure S2**: Sequence alignments of Class C (a) and subclass B2 (b) β-lactamases, highlighting sequence level conservation of amino acids. Residues highlighted in green have >85% conservation across the sequences listed. Residues highlighted in orange are conserved or catalytic residues. Multi sequence alignment was performed using T-coffee.

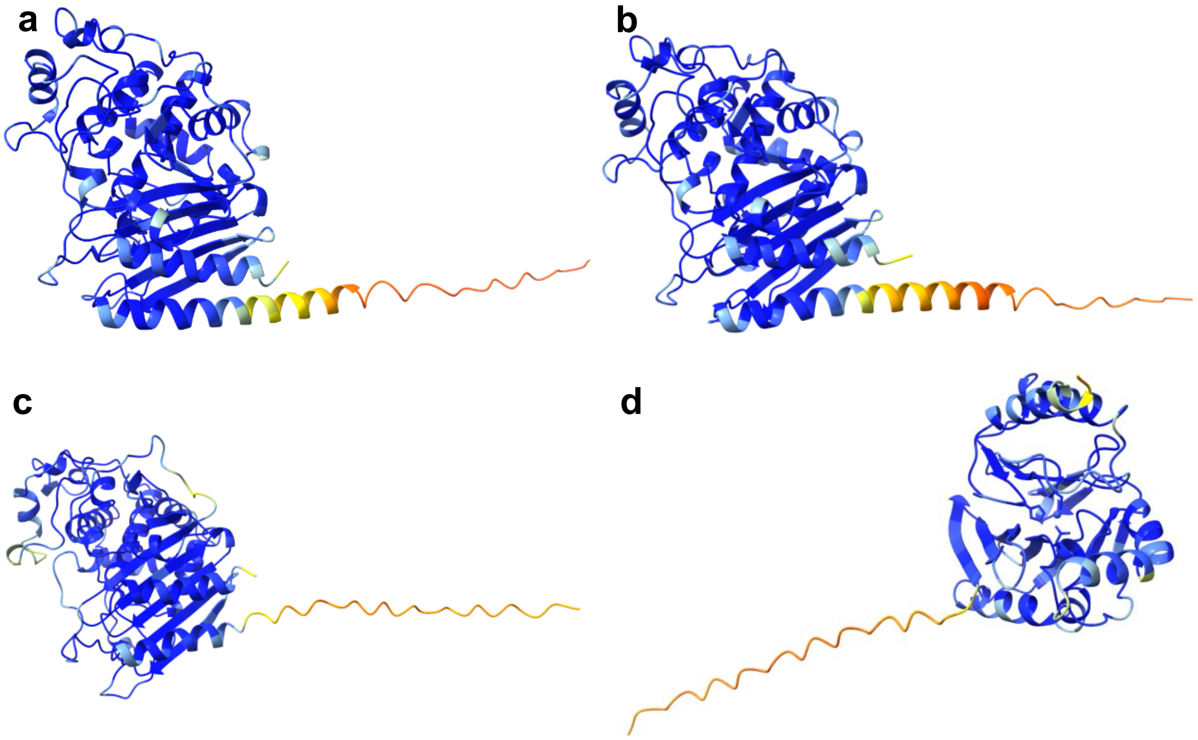


**Supplementary Figure S3**: AlphaFold predicted structures of novel β-lactamases, overlaid with colours representing the pLDDT confidence score. Dark blue = very high (pLDDT >90), Cyan = confident (90> pLDDT >70), Yellow = low (70> pLDDT >50), Orange = very low (pLDDT <50). (a) BlaPFL7. (b) BlaPFL8. (c) BlaPFL9. (d) BlaPFM5.

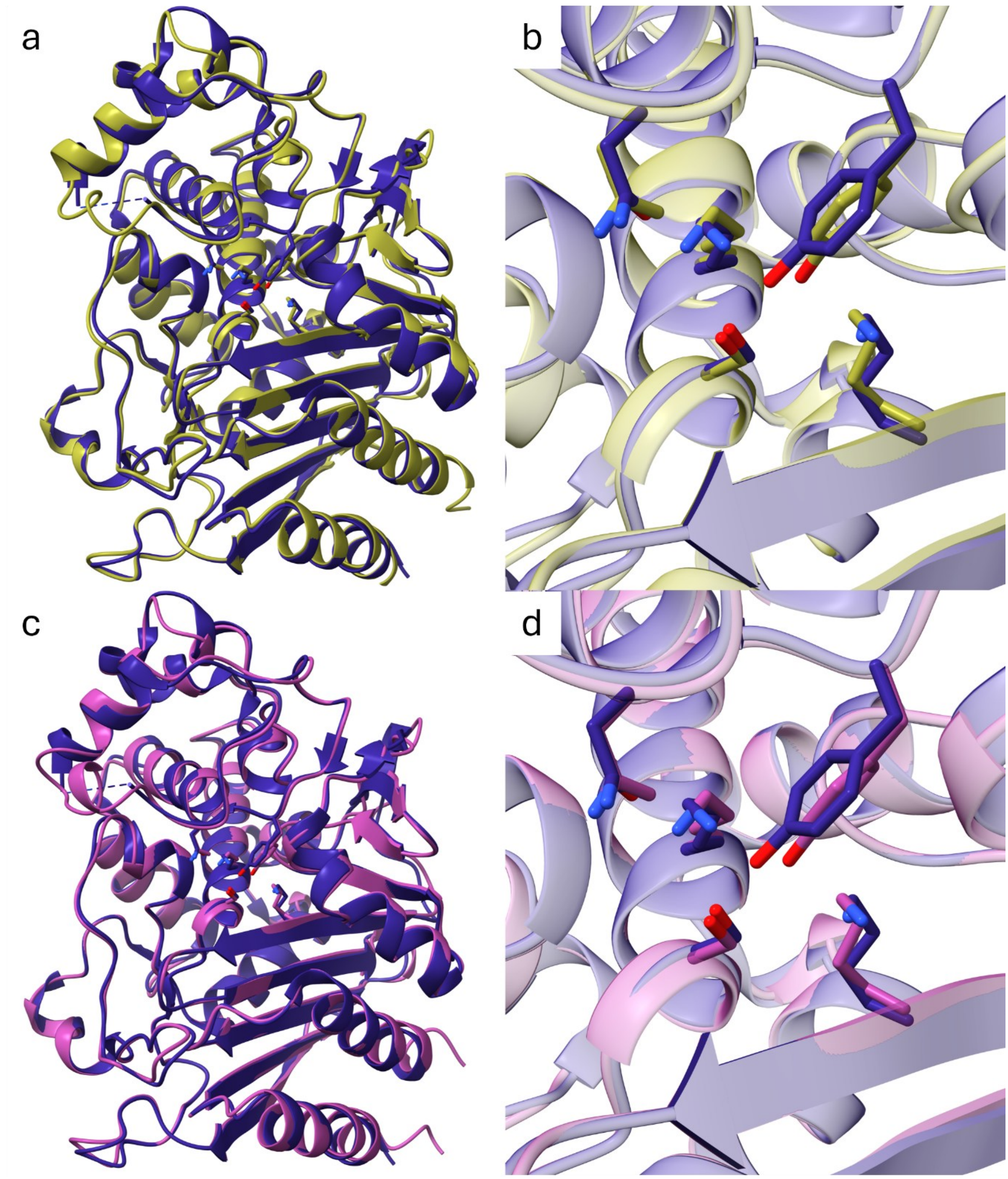


**Supplementary Figure S4**: (a) Comparison of BlaPFL8 (olive) to 2QZ6 (indigo), excluding BlaPFL8 N-terminal amino acids 1-27 for clarity. (b) Comparison of BlaPFL8 to 2QZ6 active site, highlighting the similar 3D-orientation of the catalytic residues (Ser64, Lys67, Tyr150, Asn152 and Lys315). (c) Comparison of BlaPFL9 (pink) to 2QZ6 (indigo), excluding BlaPFL9 N-terminal amino acids 1-27 for clarity. (d) Comparison of BlaPFL9 to 2QZ6 active site, highlighting the similar 3D-orientation of the catalytic residues (Ser64, Lys67, Tyr150, Asn152 and Lys315).

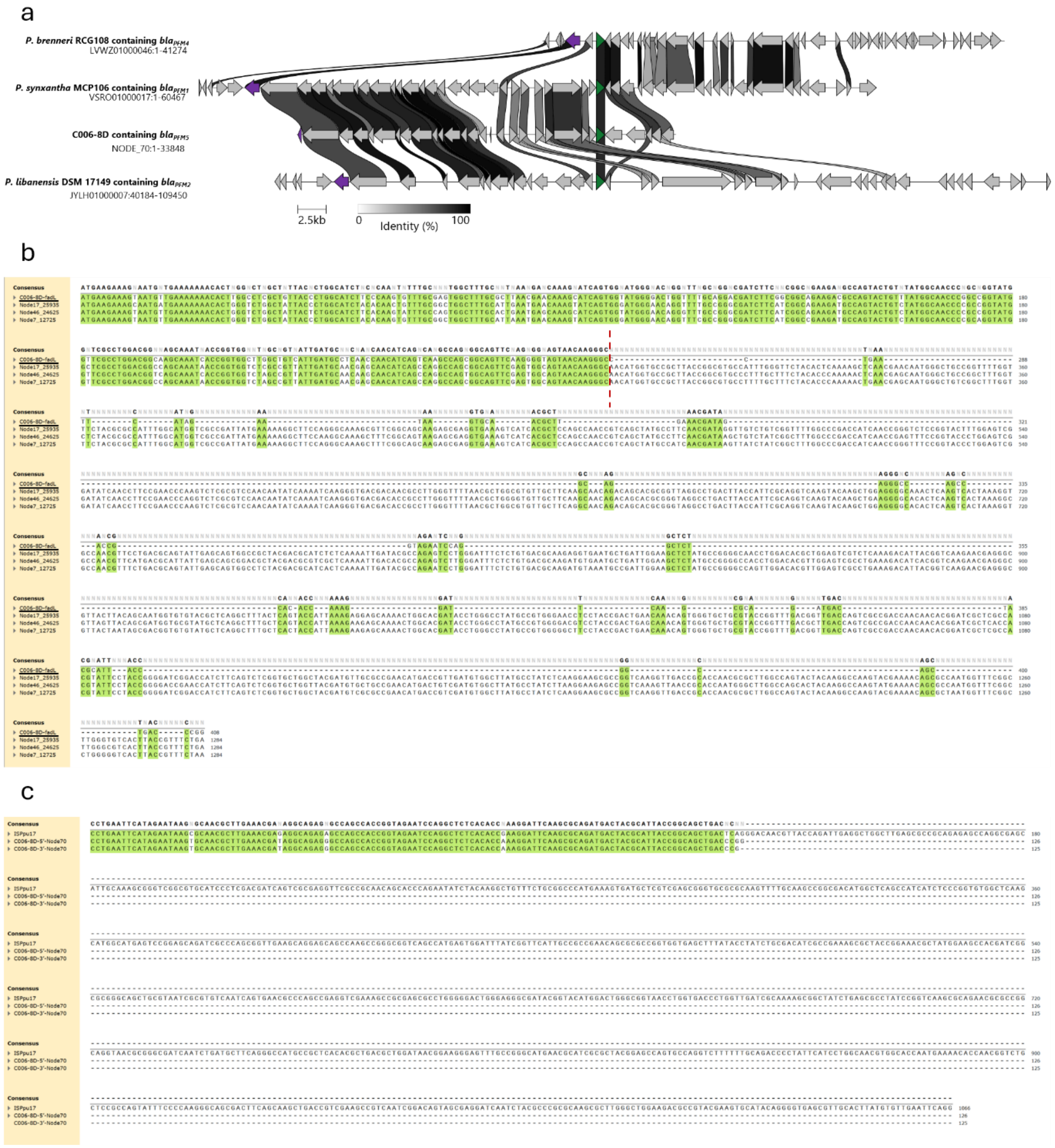


**Supplementary Figure S5**: (a) Genomic comparison of the contigs containing *bla*$_{PFM1}$, *bla*$_{PFM2}$ and *bla*$_{PFM4}$ with *bla*$_{PFM5}$, highlighting the *ompP1/fadL* transporter gene. The *bla* genes are dark green, the *ompP1/fadL* genes are purple. Figure produced using clinker [44]. (b) Sequence alignment of *ompP1/fadL* genes. Point of putative IS element insertion is indicated with a vertical dashed red. Multi sequence alignment was performed using T-coffee. (c) Sequence alignment of putative IS elements from 5' and 3' ends of contig 70 with IS*Ppu17* IS element. Multi sequence alignment was performed using MUSCLE. In (b) and (c), residues highlighted in green have >85% conservation across the sequences listed.
